# Prediction of brain age using structural magnetic resonance imaging: A comparison of accuracy and test-retest reliability of publicly available software packages

**DOI:** 10.1101/2023.01.26.525514

**Authors:** Ruben P. Dörfel, Joan M. Arenas-Gomez, Patrick M. Fisher, Melanie Ganz, Gitte M. Knudsen, Jonas Svensson, Pontus Plavén-Sigray

## Abstract

**Background:** Brain age prediction algorithms using structural magnetic resonance imaging (MRI) aim to assess the biological age of the human brain. The difference between a person’s chronological age and the estimated brain age is thought to reflect deviations from a normal aging trajectory, indicating a slower, or accelerated, biological aging process. Several pre-trained software packages for predicting brain age are publicly available. In this study, we perform a head-to-head comparison of such packages with respect to 1) predictive accuracy, 2) test-retest reliability, and 3) the ability to track age progression over time.

**Methods:** We evaluated the six brain age prediction packages: brainageR, DeepBrainNet, brainage, ENIGMA, pyment, and mccqrnn. The accuracy and test-retest reliability were assessed on MRI data from 372 healthy people aged between 18.4 and 86.2 years (mean 38.7 ± 17.5 years).

**Results:** All packages showed significant correlations between predicted brain age and chronological age (r = 0.66 to 0.97, p < 0.001), with pyment displaying the strongest correlation. The mean absolute error was between 3.56 (pyment) and 9.54 years (ENIGMA). brainageR, pyment, and mccqrnn were superior in terms of reliability (ICC values between 0.94 - 0.98), as well as predicting age progression over a longer time span.

**Conclusion:** Of the six packages, pyment and brainageR consistently showed the highest accuracy and test-retest reliability.

## 1. Introduction

Old age is the single strongest predictor for some of the most prevalent neurodegenerative disorders, such as Alzheimer’s and Parkinson’s disease [1], [2]. By accurately assessing morphological changes of the aging brain, we can potentially identify relevant pathophysiological processes, monitor disease progression, and assess the effect of neuroprotective interventions [3].

### 1.1 Brain age prediction

Patterns of region-specific changes in brain anatomy throughout the lifespan have been well described with in vivo magnetic resonance imaging (MRI) [4]. This knowledge, combined with the increasing availability of large MRI datasets, has allowed for modeling the aging brain using machine learning techniques. Such models summarize different features of the aging brain into one single measure: the predicted age, or “brain age” [5], from T1-weighted MR images. This estimate can then be used to calculate the predicted age deviation (PAD) for an individual:

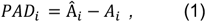

with *Â* being the predicted age and *A* the chronological age. The PAD describes the divergence from the expected aging trajectory of a brain [5], [6]. A positive PAD indicates that a brain is biologically older than what would be expected from its chronological age, which in turn could translate into an increased risk of negative age-related health outcomes. Over the past decade, several different MRI-based brain age prediction models have shown that a high PAD is predictive of developing neurodegenerative disorders, cognitive decline, and overall mortality [7]–[11].

### 1.2 Objectives

The field of brain age estimation based on MRI is fast developing, and research groups are publicly sharing pre-trained age prediction models that can be applied to new data without requiring any further training. These models range from classical regression, e.g., ridge regression [9] or Gaussian process regression [6], to gradient tree boosting [10], and more complex deep learning estimators [7], [12]–[16]. Although some studies evaluate the accuracy and reliability of brain age estimation models [17]–[19] and the use of different input features (i.e., region or voxel-based) [20], [21], no comprehensive head-to-head comparison of publicly available pre-trained models have been published to date. Such a comparison can provide an informative guide for future research that aims to implement brain age prediction software in clinical studies, e.g., to monitor age-related disease progression or the effect of interventions.

Here we compared 1) accuracy, 2) test-retest reliability, and 3) the ability to predict age progression over time for six different publicly available brain age prediction software packages with pre-trained weights.

## 2. Methods

### 2.1 Inclusion Criteria

Software packages aiming to estimate the brain age from T1-weighted MR images were included according to a set of criteria: 1) the package had to be publicly available; 2) the package had to include pre-trained weights, allowing for direct application to an independent dataset; and 3) the required pipeline for any necessary preprocessing steps of MR images had to be publically available. The following packages were identified and evaluated: brainageR [6], [22], DeepBrainNet [7], brainage [10], ENIGMA [9], pyment [13], and mccqrnn [16]. A comparison of these packages is presented in Table 1, providing an overview of the applied machine learning algorithm, input features, demographics of the training data, and reported accuracy. A more detailed explanation of the literature search is given in the supplementary material S1, where excluded packages also are listed. The package release and/or version used for the implemented packages in this study are listed in supplementary Table S1.

**Table 1.** Comparison of implemented software packages for brain age prediction.

| Package | Algorithm | Features | Original Training N, age range | Reported accuracy |
| --- | --- | --- | --- | --- |
| brainageR | GPR | voxel based (segmented) | N = 3377, 18-92 years | MAE = 4.9 years, r = 0.947 |
| brainage | GTB | region based | N = 35474, 3-96 years | r = 0.92 / 0.93 (Male/Female) |
| DeepBrainNet | 2D CNN | voxel based (T1) | N = 11729, 3-95 years | MAE = 4.12 years |
| ENIGMA | RR | region based | N = 2188, 18-75 years | MAE = 6.5 / 6.84 years<br>r = 0.854 / 0.85 years<br>(Male / Female) |
| pyment | SFCN | voxel based (T1) | N = 53542, 3-85 years | MAE = 3.9 years<br>r = 0.975 |
| mccqrnn | MCCQRNN | voxel based (segmented) | N = 10691, 20-72 years | MAE = 2.94 years |

**Model:** GPR - Gaussian Process Regression, GTB - Gradient Tree Boosting, CNN - Convolutional Neural Network, RR - Ridge Regression, SFCN - Simple Fully Convolutional Neural Network, MCCQRNN - Monte Carlo dropout Composite Quantile Regression Neural Network; **Accuracy:** MAE - Mean Absolute Error, r - Pearson's r between chronological and predicted age.

Only software packages published prior to February 2023 were included.

#### 2.1.1 brainageR

The brainageR (https://github.com/james-cole/brainageR) software package is based on Gaussian process regression, which is a non-parametric Bayesian regression approach. Briefly, the package takes raw T1-weighted MR scans as input, utilizes SPM12 (https://www.fil.ion.ucl.ac.uk/spm/software/spm12/) for segmentation into gray matter, white matter, and CSF probability maps, and calculates spatially normalized parameters. Subsequently, these normalized probability maps are concatenated into one vector and a principal component analysis is applied. The first 435 components are then used for predicting brain age with a Gaussian progress regression model. The brainageR package is implemented in R (https://www.r-project.org) and provides shell scripts for streamlined preprocessing and age prediction.

#### 2.1.2 DeepBrainNet

The DeepBrainNet (https://github.com/vishnubashyam/DeepBrainNet) package uses a two-dimensional convolutional network architecture built on top of a model trained on ImageNet, an all-purpose image database used to pre-train large deep learning models [23]. The preprocessing is fully automated and implemented in the ANTs library (https://antsx.github.io/ANTsPyNet/docs/build/html/utilities.html). The preprocessing consists of n4 bias correction, skull-stripping, and an affine registration to MNI space. Both, the package, and its implementation into the ANTs library are implemented in Python (https://www.python.org).

#### 2.1.3 brainage

The brainage (https://github.com/tobias-kaufmann/brainage) package is implemented in R and utilizes gradient tree boosting for age estimation. It uses measures of cortical thickness, area, and volume features for 180 regions of interest (ROIs) defined by the Glaser atlas [24]. FreeSurfer [25] is used to parcellate these specified cortical regions. Additionally, standard summary statistics from the FreeSurfer recon-all pipeline are used, resulting in a total of 1118 input features (i.e., 360 cortical thickness, 360 cortical area, 360 cortical volume and 38 cerebellar–subcortical and cortical summary statistics).

#### 2.1.4 ENIGMA

The package provided by the ENIGMA group applies ridge regression to predict brain age based on FreeSurfer outputs, i.e., subcortical volumes and cortical thickness, volume, and surface areas. The derived measures are averaged across hemispheres, resulting in a total of 77 features. The ENIGMA package provides separate models for males and females. It is available as a web application (https://photon-ai.com/enigma_brainage).

#### 2.1.5 pyment

The pyment package (https://github.com/estenhl/pyment-public) is based on the “Simply Fully Convolutional Network” [14], which is a 3D convolutional neural network. The model predicts brain age based on skull-stripped, MNI152 registered images. FreeSurfer is used for skull stripping, and FSL [26] is used for reorientation and linear registration to MNI space. Pyment can be used at a command line interface, docker, or application programming interface.

#### 2.1.6 mccqrnn

The package mccqrnn (https://github.com/wwu-mmll/mccqrnn_docker) implements a Monte Carlo dropout composite quantile regression neural network that adjusts for uncertainty introduced by noise in the data and the model itself. The preprocessing uses the default segmentation pipelines from the SPM12 toolbox CAT12 (https://neuro-jena.github.io/cat/). The package takes the voxels of the gray-matter segmentation as input. These are vectorized, standardized, and then used as input to the mccqrnn model. The model is implemented in Python and was trained, validated, and distributed via the PHOTON-AI (https://photon-ai.com/) software. The package can be used via Docker or as script.

### 2.2 Neuroimaging Datasets

#### 2.2.1 Subjects and study design

In total, we used data from 372 healthy volunteers, of which a subset had participated in more than one MR examination (see Table 2 and supplementary Tables S2.1, S2.2). All subject data were retrieved from the CIMBI database [27]. All participants were “healthy controls” as defined by study-specific criteria. Generally, a history or present state of neurological or psychiatric disorder was an exclusion criterion. All subjects also completed a psychical and neurological examination and a biochemical blood screening.

**Table 2:**
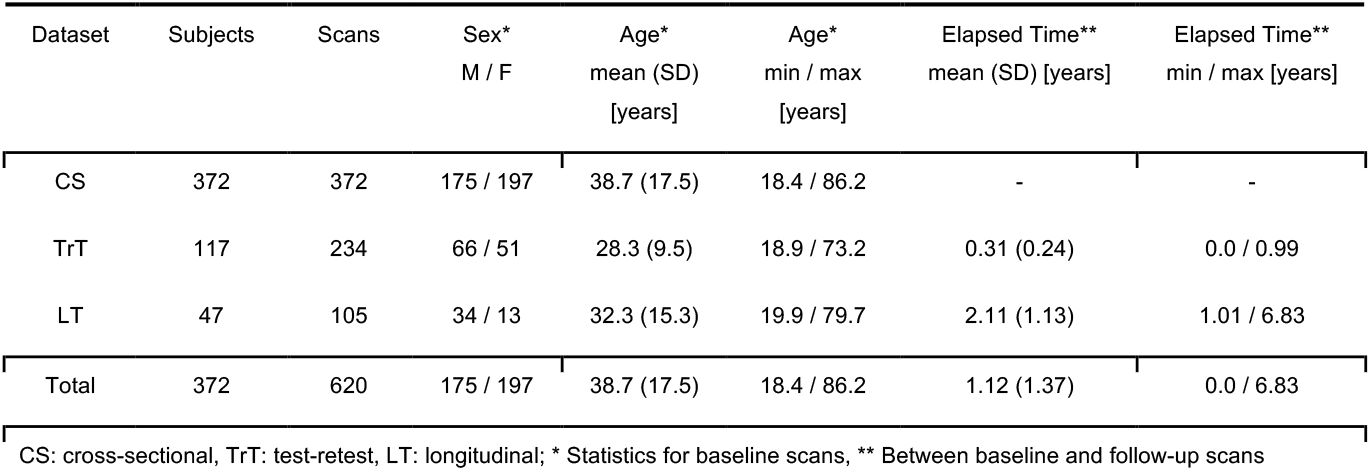
Description of included datasets.

| Dataset | Subjects | Scans | Sex*<br>M / F | Age*<br>mean (SD)<br>[years] | Age*<br>min / max<br>[years] | Elapsed Time**<br>mean (SD) [years] | Elapsed Time**<br>min / max [years] |
| --- | --- | --- | --- | --- | --- | --- | --- |
| CS | 372 | 372 | 175 / 197 | 38.7 (17.5) | 18.4 / 86.2 | - | - |
| TrT | 117 | 234 | 66 / 51 | 28.3 (9.5) | 18.9 / 73.2 | 0.31 (0.24) | 0.0 / 0.99 |
| LT | 47 | 105 | 34 / 13 | 32.3 (15.3) | 19.9 / 79.7 | 2.11 (1.13) | 1.01 / 6.83 |
| Total | 372 | 620 | 175 / 197 | 38.7 (17.5) | 18.4 / 86.2 | 1.12 (1.37) | 0.0 / 6.83 |

CS: cross-sectional, TrT: test-retest, LT: longitudinal; \* Statistics for baseline scans, \*\* Between baseline and follow-up scans

To compare 1) the accuracy, 2) test-retest reliability, and 3) the ability to track age progression over time for the brain age estimation packages, three datasets were derived from CIMBI: 1) 372 subjects between 18-86 years with at least one magnetization-prepared rapid gradient-echo high-resolution (MPRAGE) T1-weighted structural scan were identified. The first acquired, i.e., baseline, T1-weighted scans were used for a cross-sectional analysis to evaluate the accuracy of the methods; 2) 117 subjects with a follow-up scan within one year of the respective baseline were selected for a test-retest analysis to quantify the reliability of the methods; 3) 47 subjects with one or more follow-up scans more than one year after their baseline scan were included for longitudinal analysis to compare age progression in chronological and biological age (Table 2). One year was chosen as the cut-off for the test-retest analysis as it presents a trade-off between including a reasonable number of scans and minimizing the confound of actual brain aging over time. The same TrT-analysis was also performed on a subset of subjects with fewer than 31 days, 14 days, and one day between scans, respectively, to investigate the sensitivity of the TrT metrics.

#### 2.2.2 MRI Acquisition

T1-weighted MRI scans were acquired on seven different scanners, using standard parameters. Further information on MRI acquisition can be found in Knudsen et al. [27]. More information on the scanners and sequences used can be found the supplementary Tables S2.1 and S2.2.

#### 2.2.3 Quality Control

All scans passed a set of quality control measures. First, the tool MRIQC [28] was used to provide summary reports for each raw T1 image. These reports provide quality measures, such as Dietrich’s signal to noise ratio [29], and the entropy focus criterion as an indicator for blurring and motion [30]. Such quality measures were compared for all scans using boxplots. If a scan was consistently outside 1.5 times the interquartile range, it was flagged as an outlier. Flagged outliers were manually inspected and discarded if the underlying data, or processing of data, was found to be corrupted. A similar procedure was performed for the FreeSurfer reconstruction, using QATools (https://surfer.nmr.mgh.harvard.edu/fswiki/QATools) and the ENIGMA protocol (https://enigma.ini.usc.edu/protocols/imaging-protocols/) for quality control. The quality control led to the exclusion of two MR images during the first stage (raw), and seven MR images, during the second stage (FreeSurfer).

### 2.3 Statistical analysis

#### 2.3.1 Age-prediction accuracy

A cross-sectional analysis was performed on the 372 baseline scans, using the mean absolute error (MAE),

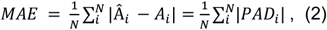

as a measure of accuracy. The MAE reflects the average absolute difference between predicted age Â_*i*_ and chronological age *A*_*i*_. For healthy controls, the MAE should, in theory, be close to zero since a healthy brain on a “normal” aging trajectory should reflect the chronological age of the individual.

Pearson’s r was calculated to assess the correlation between predicted and chronological age, and between predicted age and PAD. If the evaluated brain age model predicts chronological age perfectly, then we would observe a Pearson’s r = 1, whereas, in an unbiased model, the Pearson’s r for chronological age and PAD should be 0.

#### 2.3.2 Test-retest reliability

Test-retest reliability was evaluated using the intraclass correlation coefficient (ICC) (eq 3), specifically the ICC(1,1) [31], and the mean absolute difference (MAD) (eq 4).

The ICC can be interpreted as the proportion of variance between subjects that is due to the true signal to the total signal (true + error), and can be seen as a metrics’ ability to differentiate between subjects. A high ICC value (i.e., close to 1) means that the outcome measure can reliably rank subjects’ brain age (or PAD) estimates, a pre-requisite for doing correlation analyses. A low ICC (close to 0) means that the ranking is unreliable, and a correlational analysis using such an outcome would likely be uninformative. Specifically, ICC-1,1 is defined as follows:

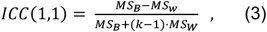

with *MS*_*B*_ and *MS*_*W*_ be the between and within subjects mean sum of squared variances, and *k* the number of observations per subject. The ICC was calculated both for brain-age predictions and the PAD values. Following recommendations from Portney and Watkins [32], the reliability will be considered to be poor when ICC < 0.5, moderate for 0.5 - 0.75, good for 0.75 - 0.9 and excellent for ICC values > 0.9.

The MAD can be seen as an estimate of the error between baseline and follow-up scan, and should be close to zero, assuming there is no measurement error or change in age-related biology. It is calculated as the absolute difference between two following brain-age estimations,

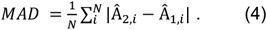

To adjust for progressed chronological age between measurements, we introduce the age adjusted MAD

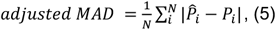

as the mean absolute difference between progressed brain age (eq 6) and progressed chronological age (eq 7),

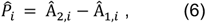

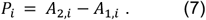

The subscripts 1 and 2 indicate baseline and follow-up scan, respectively. Here, a perfect reliability is indicated by a MAD and adjusted MAD = 0.

#### 2.3.3 Tracking age progression over time

To assess the ability of different packages to track chronological age progression, we compared the adjusted MAD (eq 5) for baseline and follow-up scans that were acquired more than one year apart. If multiple follow-up scans were available, the last follow-up scan was used. Evaluating the adjusted MAD over a longer period allows us to investigate if progressed brain-age (eq 6) corresponds to actual progressed chronological age (eq 7). A longitudinal adjusted MAD close to zero would therefore indicate that the respective package is able to accurately track chronological age over time.

To further assess the association between predicted age and chronological age, we used a linear mixed effects model, with predicted progressed biological age as dependent variable, progressed chronological age as independent variable and subjects as a random effect (to account for the subset of subjects with more than two scans). A slope-estimate close to 1 indicates that the progressed predicted age follows the actual progressed chronological age without any bias.

## 3. Results

### 3.1 Age-prediction accuracy

The results from the cross-sectional analysis based on 372 baseline scans of healthy controls are summarized in Table 3 and Figure 1. The packages showed MAE values between 3.56 years (pyment) to 9.54 years (ENIGMA).

**Table 3:** Results of Cross-Sectional, Test-Retest and Longitudinal analysis.

|  | Cross-Sectional |  |  | Test-Retest (<1 year) |  |  |  | Longitudinal (>1year) |  |
| --- | --- | --- | --- | --- | --- | --- | --- | --- | --- |
|  | MAE [years] | r | r PAD | MAD [years] | adj MAD [years] | ICC (95% CI) | ICC PAD (95% CI) | adj MAD [years] | Beta (95 % CI) |
| brainageR | 4.04 | 0.96** | -0.13* | 1.27 | 1.2 | 0.98<br>(0.98 0.99) | 0.94<br>(0.92 0.96) | 1.82 | 1.17<br>(0.68, 1.67) |
| DeepBrainNet | 6.13 | 0.89** | -0.43** | 4.25 | 4.24 | 0.57<br>(0.44 0.68) | 0.25<br>(0.08 0.41) | 4.93 | 1.93<br>(0.8, 3.05) |
| brainage | 6.39 | 0.89** | -0.59** | 2.32 | 2.35 | 0.94<br>(0.91 0.96) | 0.91<br>(0.87 0.94) | 2.88 | 0.65<br>(-0.47, 1.77) |
| ENIGMA | 9.54 | 0.66** | -0.59** | 6.05 | 6 | 0.65<br>(0.53 0.74) | 0.69<br>(0.58 0.77) | 6.49 | 1.47<br>(-1.12, 4.07) |
| pyment | 3.56 | 0.97** | -0.31** | 1.18 | 1.17 | 0.98<br>(0.98 0.99) | 0.94<br>(0.91 0.96) | 1.7 | 0.84<br>(0.44, 1.24) |
| mccqrnn | 4.46 | 0.95** | -0.46** | 1.76 | 1.73 | 0.97<br>(0.96 0.98) | 0.92<br>(0.89 0.94) | 1.98 | 1.14<br>(0.60 1.67) |

r = Pearson's correlation coefficient, associated p-value \*p<0.01, \*\*p<0.005, Beta = slope of linear mixed effects model fitted to assess relation between progressed predicted age and progressed chronological age; adj = adjusted.

**Figure 1.**
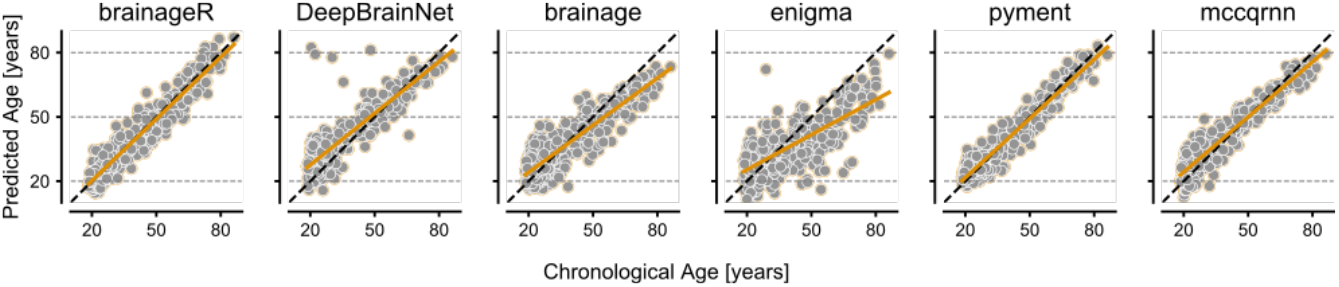
Predicted age vs. chronological age of brainageR, DeepBrainNet, brainage, ENIGMA, pyment, and mccqrnn for age-prediction on the cross-sectional dataset based on 372 scans. The identity line is in dashed black, and the model regression line is in orange.

The correlation between age and predicted age ranged from r=0.66 to r=0.97 for the packages, with the strongest association shown for pyment and the weakest for ENIGMA. Packages also showed a large range for correlations between age and PAD, r = -0.13 (brainageR) to r = -0.59 (ENIGMA and DeepBrainNet; Table 3).

### 3.2 Test-retest reliability

Test-retest results based on 117 baseline/follow-up scan pairs with less than a year elapsed between examinations are presented in Table 3. The MAD indicated that pyment (MAD = 1.18) produced the lowest deviation in estimated brain age between the two scans, while ENIGMA showed the highest deviation (MAD = 9.54). Further, the adjusted MAD was slightly lower, but very close to the unadjusted MAD, indicating that little to no age-related effects could be detected within the test-retest dataset.

The ICC showed excellent reliability for pyment, brainageR, and mccqrnn brain-age predictions and PAD values, whereas DeepBrainNet and ENIGMA showed poor to moderate reliability (Table 3). For DeepBrainNet, the poor reliability was likely driven by some extreme PAD values, which can be identified in Figure 1. A sensitivity analysis on smaller subsets for TrT analysis with baseline and follow-up scans closer in time showed a higher ICC for DeepBrainNet outcomes (Table S3.1). The ICC for PAD values followed a similar rank order as the ICC on brain-age, but with consistently lower estimates (Table 3).

### 3.3 Tracking age progression over time

The ability of the different packages to predict age progression over time was assessed on 47 subjects using their baseline and last follow-up scans (>1 year apart). The deviation between predicted brain-age progression and chronological age progression over a longer period reflects the ability of the package to accurately model the structural changes of the brain over time. The results for each method are visualized in Figure 2A and summarized in Table 3. The adjusted MAD indicates that pyment, brainageR, and mccqrnn most closely tracked age progression. The full association between progressed predicted age and progressed chronological age is depicted in Figure 2B. This analysis included any additional MR scans that subjects had participated in between their baseline and latest follow-up scan, resulting in a total of 105 scans. The slope of the fitted linear mixed effects model indicated a relationship close to 1 between predicted age progression and chronological age progression for mccqrnn, brainageR and pyment (Table 3).

**Figure 2:**
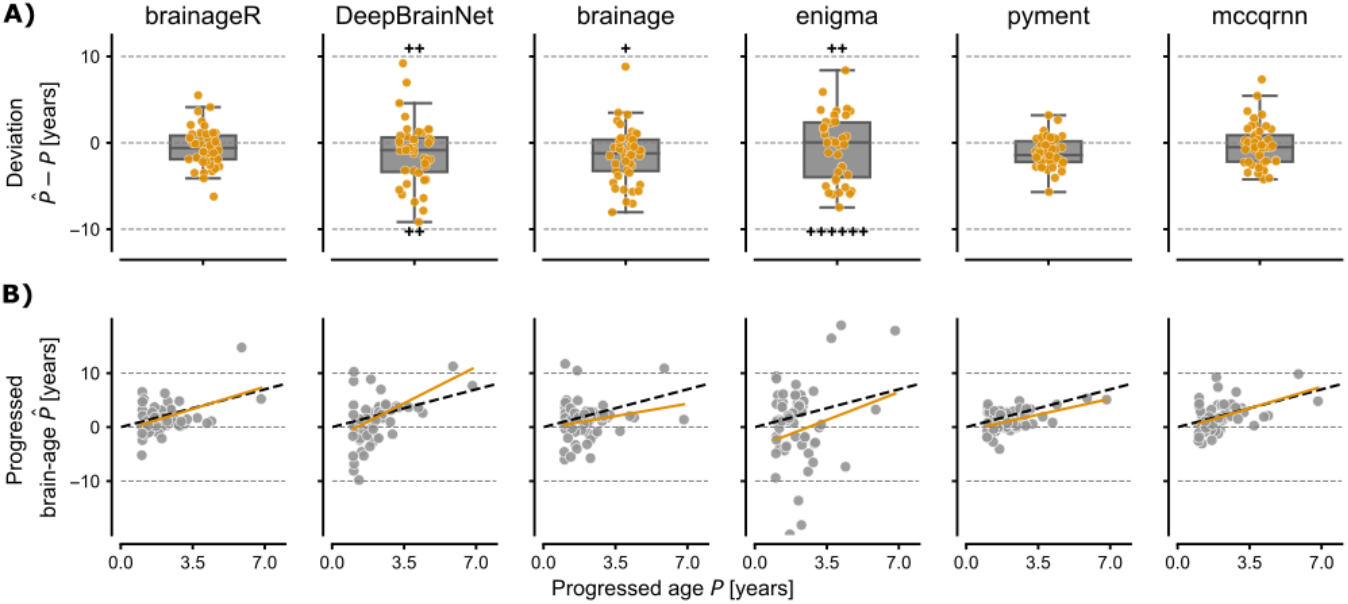
**A)** Deviation between progressed brain-age and the progressed chronological age for subjects with more than one year between MR scans. The boxplot represents the median and quartiles of the data, and the whiskers 1.5x the interquartile range. For visualization purposes, some outliers are not shown, but are instead represented by the plus-sign to indicate their quantity and respective direction. **B)** Association between chronological progressed age and progressed predicted age. The dashed black line represents the identity line, and the yellow line the fitted regression line from the LME.

## 4. Discussion

The aim of this study was to compare the accuracy and reliability of publicly available software packages for brain age prediction. We identified and applied six software packages on three datasets retrieved from the CIMBI database (Table 2) [27], data which not was used for training any of the evaluated algorithms. Out of the six software packages, pyment and brainageR showed superior performance in accuracy, test-retest reliability, and ability to predict progressed age over a longer period of time.

For brainage, brainageR and pyment, we were able to approximately reproduce the originally reported accuracies in terms of MAE and correlation between predicted age and chronological age [6], [10], [13]. However, for the packages DeepBrainNet, Enigma, and mccqrnn, we found a larger deviation from previously reported outcomes [7], [9], [16]. The three CIMBI datasets used in this study, came from several different 1.5T and 3T MR scanners and were acquired using different MPRAGE sequences (see Table S2). This heterogeneity could potentially be one reason for the discrepancy between previously reported values to those observed in this study. However, such heterogeneity can be viewed as a robustness check of the evaluated packages, as it represents real and common hardware/software heterogeneity of many existing MR datasets. As such, the results presented here have relevant construct validity with regards to the robustness of packages when confronted with such differences in data collection.

The test-retest ICC values for pyment, mccqrnn and brainageR were above 0.9 for predicted age and PAD estimates, indicating that these packages are excellent in differentiating between subjects. Such test-retest reliability is in line with previous studies, which have demonstrated high between-scan reliability of packages [8], [33]. The ICC estimates for PAD were similar to those calculated on brain-age, but consistently lower. This is to be expected, since the ICC depends on the error between scans, but also the between-subject variance. Hence, the decreased reliability for PAD values were likely due to the differences in value ranges (e.g., for pyment, the predicted biological age range is 16.5 to 68.9 years, compared to -12.0 to 10.6 years for PAD values).

A high ICC has implications for future studies of association to e.g., clinical outcomes, since a reliable ranking of subjects (based their predicted age or PAD) is a prerequisite for doing correlational analyses. The lower ICC value shown for DeepBrainNet PAD estimates suggest that a meaningful ranking of subjects is difficult to achieve with this package. Hence, an observed correlation between PAD values from this package and, e.g., a clinical outcome would likely be attenuated compared to a true, underlying association. Further, the better performing packages, brainageR and pyment, were able to reliably predict age progression over a longer time (>1 year). This is a pre-requisite for the ability to detect an effect of slowed brain aging e.g., when running a clinical trial to evaluate a putative neuroprotective drug.

It is important to note that high accuracy in predicting chronological age (i.e., MAE close to zero), does not in itself mean that the outcomes have high validity. The point of a brain age prediction is to reflect the biological state of the brain, meaning that a high deviation from chronological age can contain important information about an accelerated, or slowed, biological aging process. In a healthy population like the one used in this study, it is reasonable to assume that the MAE should be low, but a package that shows a perfect correlation to chronological age (i.e., r = 1) is unlikely to reflect information on the biological state of the brain. In line with this, it has been suggested that more loosely fitting models might be better in distinguishing pathological brains from healthy ones [7]. To further assess the construct validity of the models, future studies should compare their ability to predict e.g., negative health outcomes.

Consistent with what has previously been reported for the six software packages [6], [7], [9], [10], [13], [16], we found a negative correlation between PAD and chronological age (Table 3). Such an association is a well-known phenomenon in MR brain age prediction software packages, and is hypothesized to be caused by regression to the mean age of the training population [34]. This effect was stronger in ENIGMA, mccqrnn and brainage, compared with pyment and brainageR. A dependency of the PAD on chronological age is a potential confounder when using brain age prediction. In order to negate such effect, it has been recommended to control for chronological age when using PAD estimates to e.g., differentiate between patient groups or predict clinical outcomes[34].

Our study is not without limitations. First, the MR images used for the test-retest and longitudinal analysis were mainly collected from a sample of younger subjects. Hence, our data are less suited to draw firm conclusions on the relative performance of packages’ ability to reliably predict brain age in an older population. Second, the cutoff at one year to divide between test-retest and the longitudinal subset was arbitrarily chosen as a compromise between including a large enough N for both analyses, but avoiding any strong biological effects of aging in the test-retest analysis. A set of sensitivity analyses only including scans acquired within a shorter time frame (<1 month, < 2 week as well as < 1 day) showed similar results to the full TrT dataset for all packages, except DeepBrainNet that displayed improved performance (see supplementary Table S3).

In this study, all weights for the applied packages come from pre-training on different datasets, and no re-training was performed. Hence, no conclusion about the usability of specific machine learning algorithms is possible based on the results presented here. It was, however, not our intention to compare the advantages and disadvantages of different algorithms, but to evaluate the performance of already available packages to be used as “off the shelves” products on a new MRI dataset for brain age prediction.

## 5. Conclusion

We applied six different publicly available age prediction software packages (DeepBrainNet, brainage, brainageR, ENIGMA, pyment, and mccqrnn) to three brain MRI datasets and compared their accuracy in predicting chronological age, test-retest reliability, and ability to predict age progression over time. The packages pyment, brainageR, and mccqrnn showed the overall best performance, suggesting that they could be useful for monitoring age-related brain biology in, e.g., clinical studies. Future studies comparing the performance of these packages in predicting age-related clinical outcomes should be done to shed further light on their clinical utility.

## Supporting information

sup_lit_search_result

## Acknowledgement

This work was funded by the Impetus Longevity Grants and The Lundbeck Foundation project BrainDrugs (grant R279-2018-1145). JS was supported by Region H (grant A7167). PPS was supported by the Swedish Research Council (grant 2021-00462). We would like to thank Mark Uhrskov Juul, who’s master thesis [35] inspired this study.

