## Supplementary_material for "Prediction of brain age using structural magnetic resonance imaging: A comparison of accuracy and test-retest reliability of publicly available software packages"

**S1 Literature Search**
GitHub and PubMed were searched using search strings including "MRI", "age prediction", “brain age”, and “age estimation”.

For PubMed, abstracts were then read trough by the first author, and articles deemed potentially relevant were downloaded. Papers were scanned for methods that provided for T1w-MRI based brain age estimation and scanned for code availability. In addition, relevant references present in the downloaded article, that did not show up in the original PubMed search, were also downloaded and checked for code availability.

For GitHub results, the results were scanned for methods that provided T1w-MRI based brain age estimation, and for availability of pre-trained weights as well as inclusion of necessary pre-processing steps. Further, only packages that were linked to a published publication were included.

Identified packages are summarized in excel sheet sup_lit_search_result.xlsx. Out of 10 identified packages, six were implemented. Reasons for exclusion were missing pre-processing steps (UKBiobank_deep_pretrain, BrainAgingNet), and no pre-trained weights (BrainAgingNet). Packages (Enigma, brainage, and mccqrnn) were however included when preprocessing was based on standard pipelines with default parameters, even if such pre-processing steps was not part of the downloaded code itself. BrainAgePredictionResNet was excluded because the provided SPM preprocessing pipeline specification did not work. U-NET-for-LocalBrainAge-prediction provided unique brain-age predictions per voxels, and we considered it out of the scope of this study to include such a model and compare it to the remaining packages, who all produced a single brain age estimate.

**Table S1** The package versions / releases from implemented packages used in this study.

| | |
|---|---|
| brainageR | https://github.com/james-cole/brainageR/releases/tag/2.1 |
| DeepBrainNet | pip install antspynet==0.1.5 |
| brainage | https://github.com/tobias-kaufmann/brainage/ Commit-hash: 51e650d109347eddb420de6242fabac0485695b4 |
| Enigma | https://photon-ai.com/enigma_brainage (no version available) |
| pyment | https://github.com/estenhl/pyment-public/releases/tag/v1.0.0 |
| mccqrnn | docker image: ghcr.io/wwu-mmll/mccqrnn_docker:main-3a78f06 |

### S2 MPRAGE - MR Acquisition

**Table S2.1:** MR Scanners

| Scanner | | | | | Subset | | | | | |
|---|---|---|---|---|---|---|---|---|---|---|
| Scanner ID | Scanner | Sequences | B0 Field Strength [T] | Head Coil Channels | CS | | TrT | | LT | |
| | | | | | Subjects | Scans | Subjects | Scans | Subjects | Scans |
| d | Vision | mpr_0 | 1.5 | 8 | 44 | 44 | - | - | 5 | 10 |
| f | Trio | mpr_1, mpr_2 | 3 | 8 | 128 | 128 | 64 | 128 | 34 | 75 |
| g | Verio | mpr_4 | 3 | 32 | 61 | 61 | 6 | 12 | - | - |
| m | Trio PET/MR | mpr_3, mpr_4, mpr_5 | 3 | 12 (43) 32 (12) | 23 | 23 | 11 | 22 | 1 | 3 |
| n | Prisma | mpr_7, mpr_8 | 3 | 32 (103), 20 (1) | 72 | 72 | 23 | 46 | 1 | 2 |
| p | Prisma Fit | mpr_6 | 3 | 64 | 41 | 41 | 13 | 26 | 6 | 15 |
| v | Verio | mpr_4 | 3 | 32 | 3 | 3 | - | - | - | - |

**Subset**: CS – Cross-Sectional, TRT- Test-Retest, LT – Longitudinal;
**Head Coil Channels**: if different receiver coils were used on a scanner, the number of scans using the specific coils were indicated in brackets i.e.: 12 (43) – 12 channels, 43 scans.

**Table S2.2:** MR acquisition parameters

| Sequence ID | TI [ms] | TE [ms] | TR [ms] | FlipAngle [°] | Scanner ID* | Subsets | | | |
|---|---|---|---|---|---|---|---|---|---|
| | | | | | | Total | CS | TrT | LT |
| mpr_0 | 100 | 4.4 | 11.4 | 8 | d | 49 | 44 | 0 | 10 |
| mpr_1 | 800 | 3.04 | 1550 | 9 | f | 155 | 103 | 86 | 27 |
| mpr_2 | 800 | 3.93 | 1540 | 9 | f | 107 | 25 | 42 | 48 |
| mpr_3 | 900 | 2.26 | 2300 | 8 | m | 12 | 4 | 8 | - |
| mpr_4 | 900 | 2.32 | 1900 | 9 | g,m,v | 107 | 72 | 12 | - |
| mpr_5 | 900 | 2.44 | 1900 | 9 | m | 20 | 11 | 14 | 3 |
| mpr_6 | 900 | 2.58 | 1900 | 9 | p | 66 | 41 | 26 | 15 |
| mpr_7 | 920 | 2.41 | 1810 | 9 | n | 11 | 4 | 3 | - |
| mpr_8 | 972 | 2.58 | 2000 | 8 | n | 93 | 68 | 43 | 2 |

*See Scanner ID in Table S2.1. **Subset**: CS – Cross-Sectional, TRT- Test-Retest, LT – Longitudinal

### S3 TrT Subset Analysis

**Table S3.1** Subset test-retest analyses with shorter between-scan intervals. MAD and ICC values are shown for the predicted brain age.

| Method | MAD [Years] | | | | ICC [CI95%] | | | |
|---|---|---|---|---|---|---|---|---|
| | <1 Day | <14 Days | <31 Days | <365 Days | <1 Day | <14 Days | <31 Days | <365 Days |
| brainageR | 0.78 | 0.94 | 1.16 | 1.27 | 1 [0.99 1.0] | 0.99 [0.98 1.0] | 0.99 [0.98 0.99] | 0.98 [0.98 0.99] |
| DeepBrainNet | 0.6 | 1.24 | 1.72 | 4.25 | 1 [1.0 1.0] | 0.98 [0.95 0.99] | 0.97 [0.94 0.98] | 0.57 [0.43 0.68] |
| brainage | 1.68 | 2.05 | 1.93 | 2.32 | 0.98 [0.9 1.0] | 0.95 [0.86 0.98] | 0.96 [0.93 0.98] | 0.94 [0.92 0.96] |
| ENIGMA | 6.58 | 5.26 | 7 | 6.05 | 0.46 [-0.26 0 .86] | 0.46 [-0.03 0 .78] | 0.36 [0.05 0.61] | 0.65 [0.53 0.74] |
| pyment | 1.04 | 0.95 | 1.01 | 1.18 | 1 [0.98 1.0] | 0.99 [0.98 1.0] | 0.99 [0.99 1.0] | 0.98 [0.98 0.99] |
| mccqrnn | 2.08 | 1.78 | 1.67 | 1.76 | 0.99 [0.93 1.0] | 0.98 [0.96 0.99] | 0.98 [0.97 0.99] | 0.97 [0.96 0.98] |

**Table S3.2** Demographics for Subset Analysis.

| Subsets | Subjects | Scans | Sex* M/F | Age* [years] Mean / SD | Age* [years] Min / max | Elapsed Time** mean / SD [years] | Elapsed Time** min / max [years] |
|---|---|---|---|---|---|---|---|
| <1 Day | 8 | 16 | 2 / 6 | 30.87 / 18.2 | 18.89 / 73.15 | - | - |
| <14 Days | 15 | 30 | 7 / 8 | 32.14 / 14.65 | 18.89 / 73.15 | 0.01 / 0.02 | 0.0 / 0.04 |
| < 31 Days | 37 | 74 | 22 / 15 | 29.51 / 11.69 | 18.89 / 73.15 | 0.04 / 0.03 | 0.0 / 0.08 |
| < 365 Days | 117 | 234 | 66 / 51 | 28.28 / 9.47 | 18.89 / 73.15 | 0.31 / 0.24 | 0.0 / 0.99 |

*Statistics for baseline scans; ** Between baseline and follow-up scans
